# recountmethylation enables flexible analysis of public blood DNA methylation array data

**DOI:** 10.1101/2022.05.19.492680

**Authors:** Sean K. Maden, Brian Walsh, Kyle Ellrott, Kasper D. Hansen, Reid F. Thompson, Abhinav Nellore

## Abstract

Thousands of DNA methylation (DNAm) array samples from human blood are publicly available on the Gene Expression Omnibus (GEO), but they remain underutilized for experiment planning, replication, and cross-study and cross-platform analyses. To facilitate these tasks, we augmented our `recountmethylation` R/Bioconductor package with 12,537 uniformly processed EPIC and HM450K blood samples on GEO as well as several new features. We subsequently used our updated package in several illustrative analyses, finding (1) study ID bias adjustment increased variation explained by biological and demographic variables, (2) most variation in autosomal DNAm was explained by genetic ancestry and CD4+ T-cell fractions, and (3) the dependence of power to detect differential methylation on sample size was similar for each of peripheral blood mononuclear cells (PBMC), whole blood, and umbilical cord blood. Finally, we used PBMC and whole blood to perform independent validations, and we recovered 40-46% of differentially methylated probes (DMPs) between sexes from two previously published epigenome-wide association studies (EWAS).

## 1 Introduction

DNA methylation (DNAm) is the most commonly studied epigenetic mark, and most public DNAm array samples are generated from blood [1]. In prior work [1], we conducted comprehensive cross-study analyses of human DNAm array studies with raw data deposited on the Gene Expression Omnibus (GEO) [2, 3], the largest archive of publicly available array data. We confined attention to the HumanMethylation450K (HM450K) platform introduced by Illumina in 2012. HM450K arrays profile 485,577 CpG loci concentrated in protein-coding genes and CpG island regions [4, 5]. We found that: (1) a subset of Illumina’s prescribed BeadArray controls explained most quality variances; (2) samples clustered by tissue and cancer status in a principal component analysis (PCA) of autosomal DNAm; and (3) subsets of CpG probes showed high tissue-specific DNAm variation among 7 normal tissues. We further released the `recountmethylation` Bioconductor package [6] along with uniformly processed data compilations pairing DNAm with harmonized metadata labels for age, sex, tissue, and disease state.

The initial `recountmethylation` release left open several important issues. First, the delay between data compilation and reporting meant our initial release of data compiled in March 2019 was over a year out of date at the time of publication. Second, the prevalence of raw data from the newer EPIC platform [7] is rapidly increasing while our initial data compilation included only samples run on the older HM450K platform. Finally, several practical research concerns were not accommodated in the initial package release, including how to leverage public array data compilations to determine the required number of samples to test a new hypothesis, how to account for confounding factors in cross-study analyses, and how to leverage public data to independently validate previously published DMPs and identify subsets of high-confidence biomarker candidates.

We address these outstanding issues in the present paper using novel cross-study and cross-platform analyses, confining attention to normal human blood samples. Blood DNAm is often probed in epigenome-wide association studies (EWAS) to discover, test, and validate biomarkers [8–10] for diseases such as type II diabetes [11–13], obesity [14], non-alcoholic fatty liver disease [15], asthma [16], and dementia [17], as well as colorectal [18–21], esophageal [22], breast [23], pancreatic [24], and head-and-neck [25] cancers. It is widely used to study biological aging [26–29] and normal tissue epigenetics [30, 31], including development and function of the immune system [32]. Recent work studied how gestational age-related differential DNAm relates to fetal health and disease risk [28, 33]. Further, cord blood DNAm is increasingly used to precisely quantify fetal gestational age [29, 34–36], which may lead to improvement in the efficacy of prenatal screening [37]. In addition, many software tools were trained and designed for use with blood DNAm data; these included methods for cell-type deconvolution [38–40], inference of population genetic structure and shared genetic ancestry [41], and power analyses [42].

DNAm differences between sexes have been observed in mouse [43] and multiple human tissues including brain [43], pancreas [44], nasal epithelium [45], cord blood [46], and whole blood [47, 48]. These DNAm differences can impact insulin secretion [44], risk of disease [45, 46], and biological age [43]. Using cross-study and cross-platform compilations of whole blood and PBMC, we performed novel independent validation of previously published sets of probes with differential methylation between the sexes (a.k.a. “sex DMPs”) from two previous studies in whole blood [47, 48].

## 2 Results

### 2.1 12,537 normal blood samples spanning 3 sample types were incorporated into recountmethylation

We uniformly processed raw intensity data generated on the HM450K or EPIC platforms for 68,758 samples available on GEO before March 31, 2021 (Fig. 1, Methods, [49, 50]). We narrowed focus to 12,537 normal human blood samples from 63 studies, each of which had *≥* 10 samples after quality control. After harmonizing metadata across studies, we found these samples were predominantly of three types (Fig. 2a): whole blood, umbilical cord blood (a.k.a. “cord blood”), and peripheral blood mononuclear cells (a.k.a. “PBMC”). Whole blood was distinguished from PBMC by the presence of erythrocyte and granulocyte DNA, as these cell types are removed during PBMC preparation [51] (Fig. 2a). Each blood sample type included *≥* 245 samples from *≥* 2 studies per respective platform (Fig. 2b, Table S1).

**Fig. 1:**
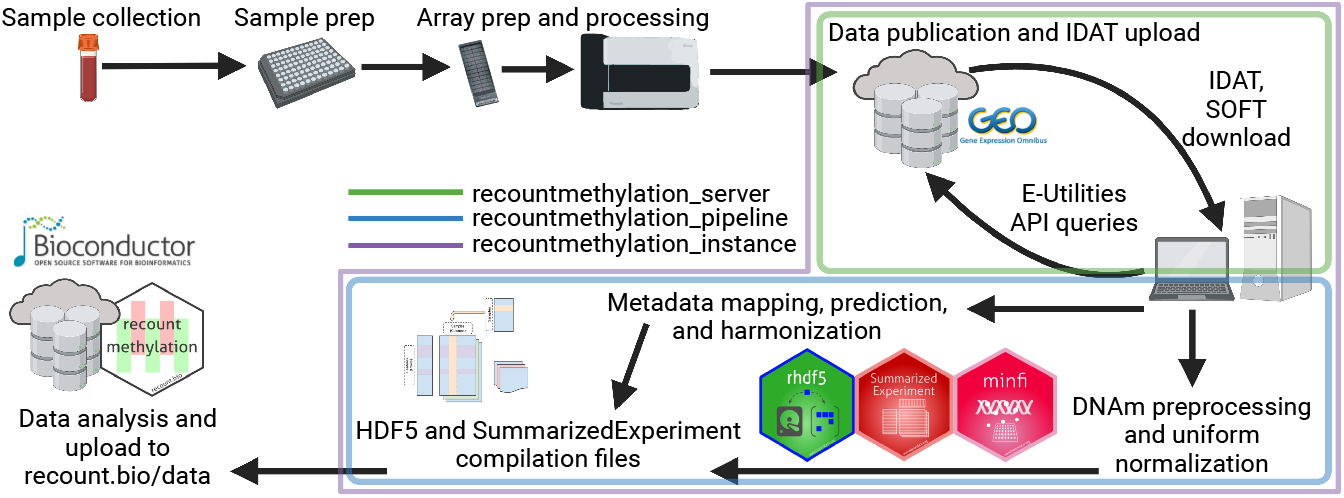
Workflow to obtain public DNAm array data from GEO. Collection, preparation, and processing of array samples (top left) as well as publication of GEO datasets were performed by other investigators (top right). We downloaded raw intensity data (IDATs) and metadata (SOFTs; top right), processed GEO metadata (middle) and DNAm signals (bottom right) into HDF5-based data formats (bottom middle), and finally updated our server and the `recountmethylation` Bioconductor package (bottom left). Color outlines indicate data access and processing using tools we developed (green = `recountmethylation server` [52], blue = `recountmethylation pipeline` [53], green = `recountmethylation instance` [54]). Diagrams were created with BioRender.com.

**Fig. 2:**
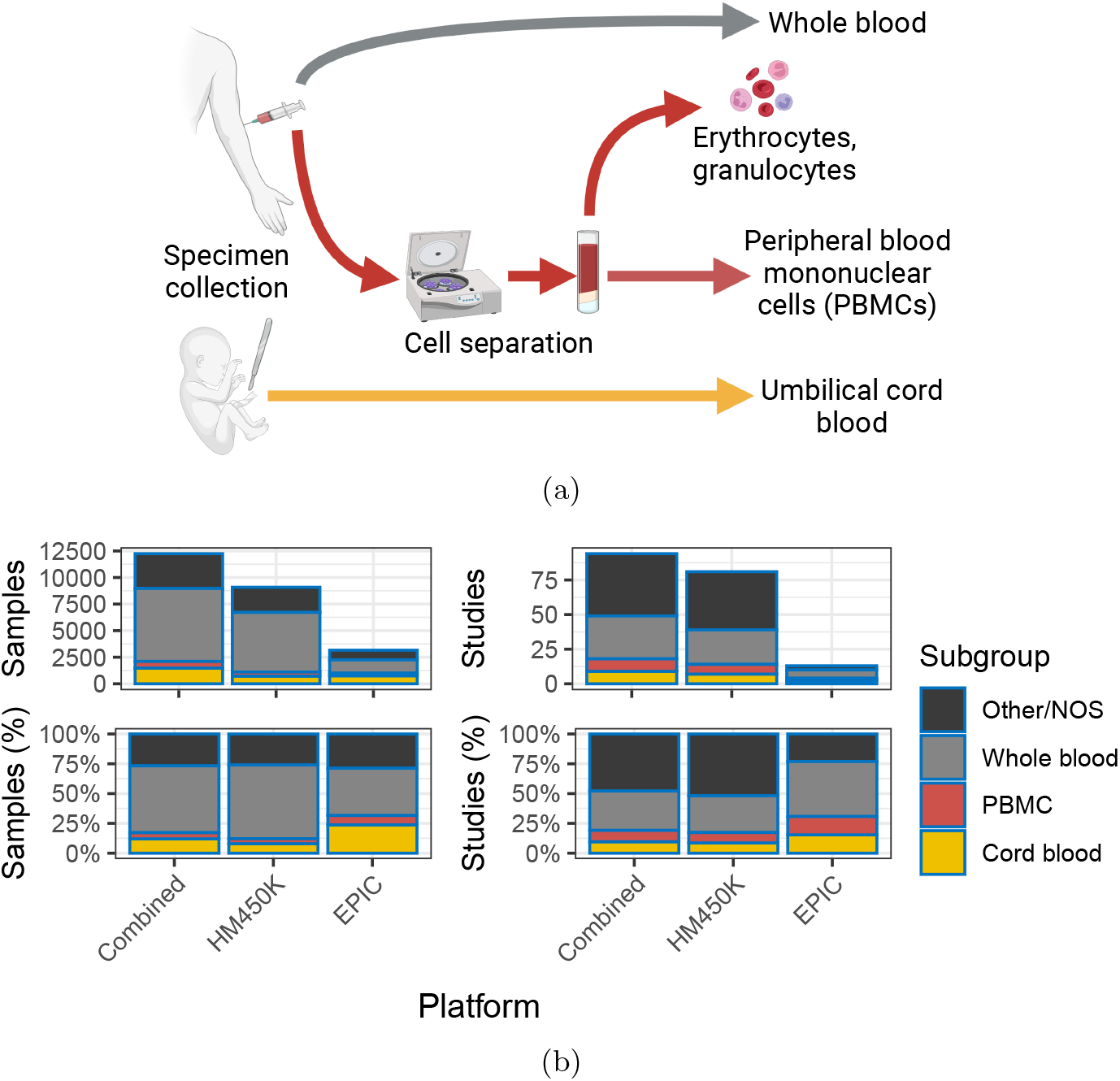
Blood specimen collection and DNAm array data availability by sample type. (a) Blood sample collection and handling prior to upload to GEO. (b) Barplot summaries of available samples (left) and studies (right), showing counts (top) and percentages (bottom) of blood sample types (black = other/not otherwise specified [NOS], gray = whole blood, red = PBMCs, yellow = cord blood). Bar heights indicate the aggregate sample group “all.” Diagrams were created with BioRender.com.

We subsequently updated our Bioconductor package `recountmethylation` [1] to facilitate cross-study and cross-platform analyses of the blood samples. The package’s new features permit search for samples with DNAm profiles similar to a query sample [55], inference of shared genetic ancestry [41], and novel power analyses [42]. These features are explained in package vignettes. Further, a new `recountmethylation instance` Snakemake workflow available on GitHub [54] allows users to create their own compilations of public DNAm array data on GEO [56], with the functionality to customize output data types and attributes predicted from GEO metadata. As shown below, our resources enable identification of biomarker candidates, independent validation and replication of previous research, experiment planning, and more.

### 2.2 Study ID adjustment increased variation explained by biological and demographic variables

We conducted simulations investigating the impact of bias correction by study ID, a surrogate for technical confounders [1]. Three DNAm values were modeled in multiple regressions: (1) unadjusted DNAm, (2) uniform adjustment on 5 randomly selected studies (a.k.a. “adjustment 1”), and (3) exact adjustment on 2-4 randomly selected studies (a.k.a. “adjustment 2”). Regression models 2 and 3 were compared to test whether two distinct study ID bias adjustment strategies had comparable outcomes. We determined the fraction of explained variance (FEV) for each of 13 variables from ANOVA, yielding 3 results per variable per simulation rep (Methods). Total non-residual variances almost invariably decreased after applying either of the 2 study ID adjustment strategies (Fig. S2a, median fractions of non-residual variances, adjusted over unadjusted, adj. 1 = 6.88e-1, adj. 2 = 6.84e-1). Variance reduction magnitudes were identical across adjustment strategies, with the exception of a few outlying models from adjustment 1 simulations (Fig. S2b).

We categorized variables as biological (e.g., six predicted blood cell type fractions), demographic (e.g., predicted sex, age, and genetic ancestry), or technical (e.g., platform, where applicable). Across all 3 variable categories, FEV increased relative to unadjusted models after either adjustment strategy, and FEV distributions were far more similar among adjusted models than between adjusted and unadjusted models (Fig. 3a). The largest median FEV differences were observed for demographic variables, while the smallest were observed for technical variables. Among individual variables, median FEV was *<* 0.1 across most models and variables, where study ID showed the maximum median FEV of 0.47 for unadjusted DNAm. After either adjustment, study median FEV decreased drastically to *≤* 2e-3, while median FEV for all remaining variables increased (Table S2).

**Fig. 3:**
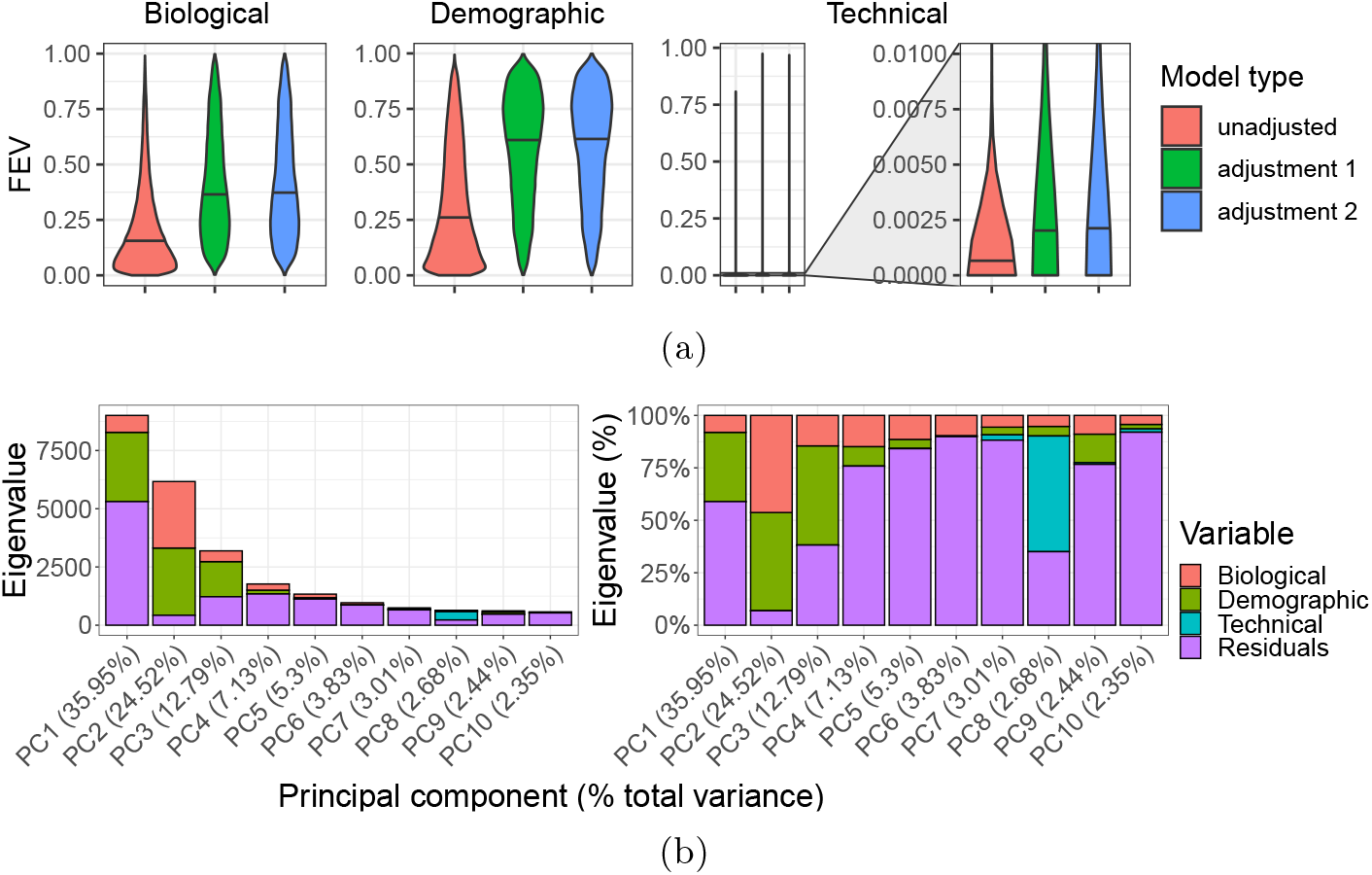
Variance analyses of study bias adjustments and principal components. (a) Distributions of fraction of explained variance (FEV). Violin plots show results grouped by 3 variable categories (plot titles, one of biological on left, demographic in middle, or technical on right), and color fills show model type (pink = adjustment 1 or adjustment on 5 studies, green = adjustment 2 or adjustment on 2-4 studies, and blue = unadjusted). (b) Autosomal DNAm principal component analysis (PCA) results across normal blood samples. Stacked barplot y axes show the eigenvalue magnitudes at left and percentages at right for the top ten components on the x axes. Fill colors indicate magnitudes of component sum of squared variances explained by variable categories (red = biological, green = demographic, blue = technical, purple = residuals).

Because performing compilation-wide corrections on study ID substantially increased variation explained by biological and demographic variables, we included Beta-values under our adjusted models in `recountmethylation` for reuse in cross-study analyses.

### 2.3 Most explained DNAm variation is from predicted genetic ancestry and predicted cell composition

To better understand key sources of variation in compiled blood data, we performed principal component analysis (PCA) on normalized [49], study ID-corrected autosomal DNAm, followed by ANOVA on regressions with 13 variables categorized as either biological, demographic, or technical (Methods). While most variation was residual across most components, explained variation was mainly from demographic variables at components 1 and 3, biological variables at components 4,5,6, and 10, and from technical variables at component 8, and split between demographic and biological variables at component 2 (Fig. 3b). Most explained variation from demographic variables was from genetic ancestry in the first component, and while CD4+ T-cell fraction explained substantial biological across remaining top components (Fig. S3). The top two principal components showed samples clustered largely independent from sample type and platform labels, but showed distinct gradient patterns for genetic ancestry, CD8+ T-cells, CD4+ T-cells, and B-cells (Fig. S4).

### 2.4 Dependence of statistical power on sample size was similar across blood sample types

We conducted power analyses on the blood samples included in `recountmethylation` by applying the simulation-based `pwrEWAS` approach [42] (Methods). To attain *≥* 80% power to detect DMPs between two groups of roughly equal size, the *N* estimated total samples required were similar across sample types, where *N ≈* 300 samples at mean Beta-value difference between groups *δ* = 0.05, *N ≈* 150 samples at *δ* = 0.1, and *N ≈* 80 samples at *δ* = 0.2. We assumed an FDR threshold of 5%. Outcomes were similar within each of the whole blood, cord blood, and PBMC groups, but they were worse when including all blood samples, likely due to greater sample heterogeneity (Fig. 4).

**Fig. 4:**
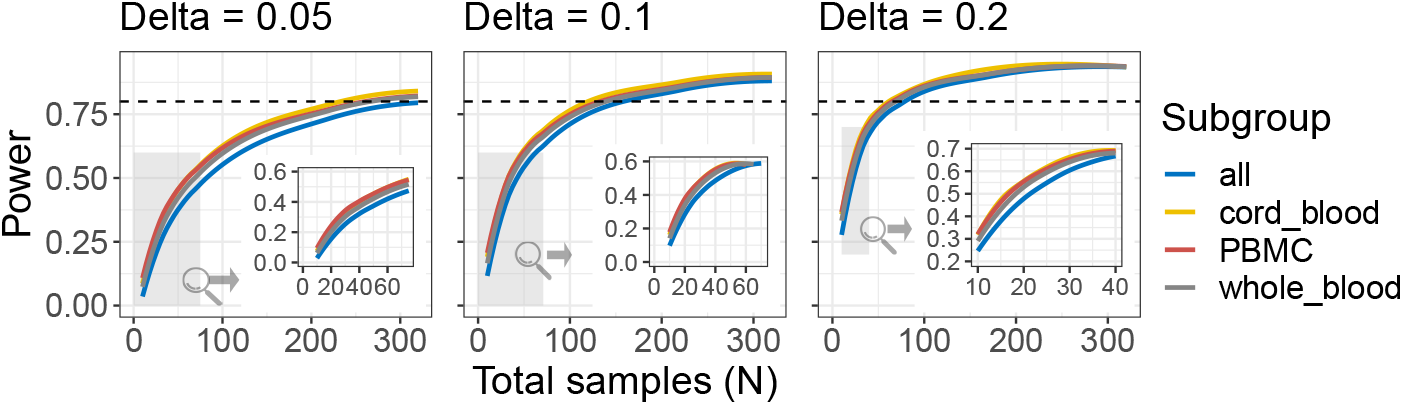
Results of power analyses with the simulation-based `pwrEWAS` method [42]. Curves indicate tradeoffs between power on the y axes and total samples *N* on the x axes, across 3 delta values (0.05 at left, 0.1 at middle, 0.2 at right). Curve colors show results for each sample type (line colors, blue = all, yellow = cord blood, red = PBMCs, gray = whole blood). Dotted horizontal lines show the 0.8 (80%) target power. Grayed regions and insets show magnifications of regions with low *N*.

Our results suggest fewer samples are necessary than the results of [42], where adult PBMCs showed *≥*80% power with *N* = 220 samples at *δ* = 0.1. Further, an independent power analysis using whole-blood EPIC arrays [57] found 85% of probes had *>*80% power with *N* = 200 and *δ* = 0.1, although their FDR cutoff value of 15% was less stringent than our cutoff value of 5%.

### 2.5 40% of sex DMPs from a previously published EWAS study were replicated in either whole blood or PBMC

We queried a search index of blood autosomal CpG DNAm, which is included in the updated `recountmethylation` resource, for each of the 113 whole blood samples from [48]. In the process, we quantified the similarity of queried sample methylation profiles to other samples by analyzing the *k* nearest neighbors returned (Methods). Among the 1,000 nearest neighbors returned per queried sample, the whole blood label was common while the PBMC label was rare (Fig. 5a), in agreement with Methods in [48] describing the queried samples as “peripheral whole blood.” This greater similarity to compiled whole blood may reflect greater similarity in subject ages, cell composition [51], and/or genetic ancestry (Fig. S5a), and we corrected for these potential confounders in regressions for identifying sex DMPs from either compilation (Methods).

**Fig. 5:**
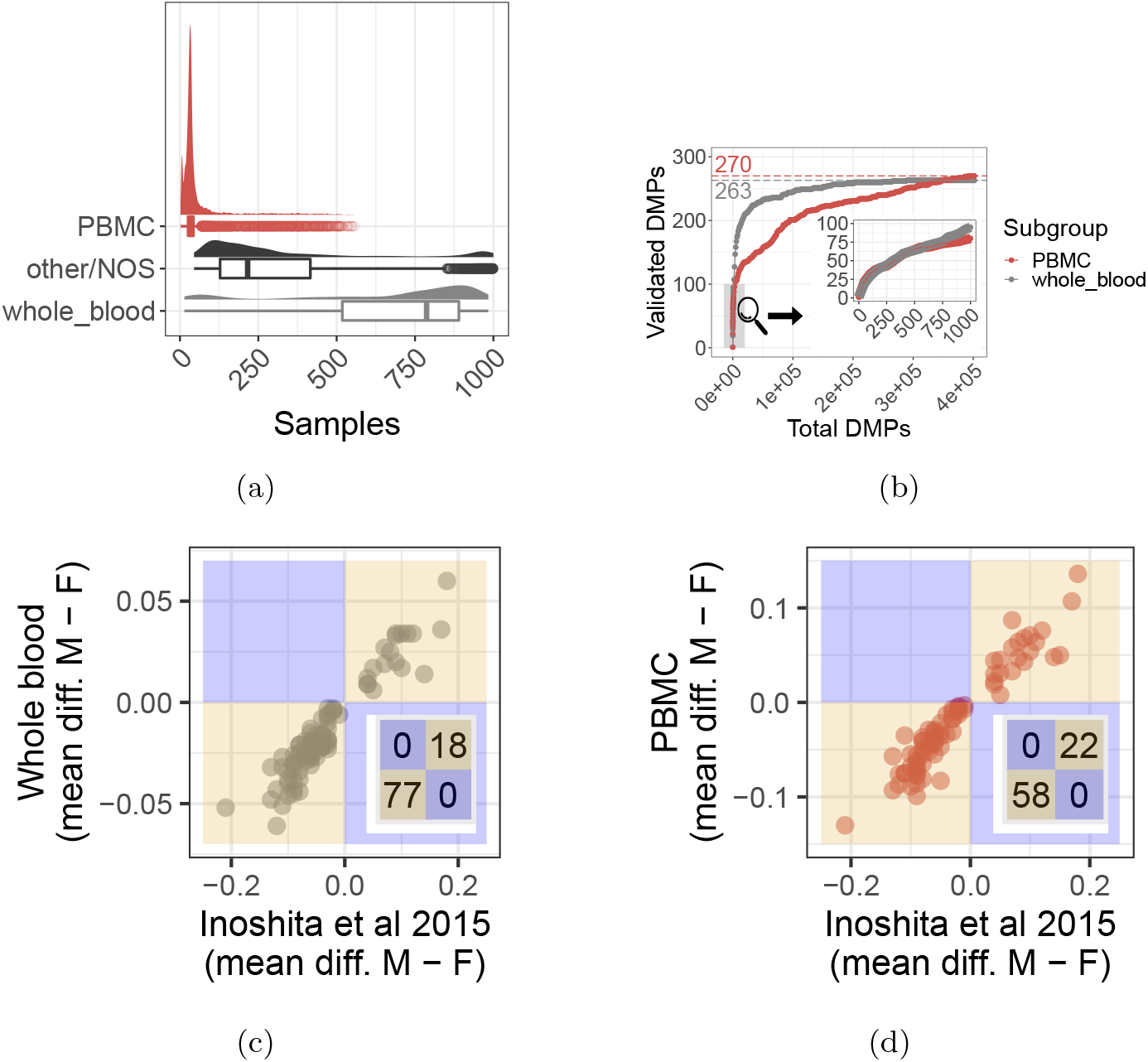
Replication of sex DMPs from [48] (a.k.a “Inoshita et al 2015”) using cross-study compilations of whole blood and PBMC. (a) Sample label distributions among the 1,000 nearest neighbors from querying [48] samples, where density and box plots show returned frequencies of 3 sample labels (red = PBMCs, black = other/not otherwise specified [NOS], gray = whole blood). (b) Concordance of [48] DMPs on the y-axis among the top significant compilation DMPs on the x-axis, ranked on P-values, for whole blood in gray and PBMCs in red. The zoom shows the top 1,000 DMPs, and colored dotted lines and colored numbers indicate total DMPs from [48] (Methods). (c and d) Mean Beta-value differences (male -female) at [48] DMPs on y axes and in (c) whole blood and (d) PBMC compilations. Region colors show direction agreement in gold and disagreement in blue, and insets show DMP counts by region.

We next considered the threshold of the top 1,000 most significant DMPs from whole blood and PBMC. We set this threshold because these DMPs captured the long tail of high between-sex DNAm differences for each tissue (Fig. S6a- S6d), and because of less replication divergence between tissues compared to less stringent thresholds observed from our concordance at the top analysis (Fig. 5b). Among these whole blood and PBMC DMPs, (117/292 =) 40% replicated sex DMPs from [48] (Fig. 5b). Of these, 95 were only replicated in whole blood, 80 were only replicated in PBMC, and 58 were replicated in both tissues (Fig. S7a). Further, (248/544 =) 46% of whole-blood sex DMPs independently reported in [47] overlapped DMPs in whole blood or PBMC. However, just 26 sex DMPs appeared in all of PBMC, whole blood, [48], and [47]. Mean (normalized) Beta-value was typically higher in females than males in whole blood (64-81% of DMPs) but not PBMC (35% of DMPs). There was high agreement in mean Beta-value differences between males and females, with slightly greater agreement among whole blood than PBMC (99% *>* 96% direction agreement) across sex DMPs from [48], and 100% agreement among the subset of replicated DMPs in each compilation (Figs 5c and 5d). DNAm proximal to cytosine- and guanine-rich regions, known as CpG islands, can functionally regulate gene expression [58, 59, 59–62], and most replicated sex DMPs mapped to CpG islands (65/117, 56%). The most significant of these DMPs mapped to a variety of gene regions, including two body DMPs at *RFTN1* and *LOC644649*, and one promoter DMP at *SLC6A4* (P-adjusted *<* 5.1e-47, Bonferroni method, Supplementary Materials).

## 3 Discussion

We analyzed DNAm array data from the three most prevalent blood sample types in the GEO database and updated the `recountmethylation` Bioconductor package to make reproducible [63, 64] cross-study and cross-platform analyses of these data easier. Since HM450K and EPIC data continue to accumulate rapidly on GEO, we further developed the `recountmethylation instance` Snakemake workflow to enable semi-automated compilation of the DNAm array data on GEO [54].

We replicated 40% of sex DMPs from [48] and 46% of sex DMPs from [47] using independent whole blood and PBMC compilations. These rates were similar to prior studies of sex DNAm differences, including a 38% validation rate of cord blood sex DMPs between two independent cohorts [46], and 44% validation rate of genes in nasal epithelium with DNAm differences by sex [45]. These results could represent a baseline expectation for replication or independent validation rate of DMPs for sex, and potentially other variables, across independent EWAS.

Our work has several limitations. First, we excluded blood spots from our analyses due to insufficient raw DNAm array data available from GEO, although this blood sample type accounts for a substantial fraction of publicly available data from younger subjects. Another limitation related to data availability is that far fewer blood samples were available for the EPIC platform compared to HM450K as of March 31, 2021. The larger EPIC platform could help expand analyses to new genome regions and clarify regional DNAm signals at CpG islands and genes. The `pwrEWAS` method assumes a technical detection threshold of Beta-value = 0.01 by default, and using this threshold ensures our findings are relevant for both single-study and cross-study analyses. However, this technical threshold likely should be lowered if the study being planned involves cross-study analyses using study ID bias correction, because we found this correction reduced explained variances (Section 2.2) and resulted in lower between-group differences in our sex DMP cross-study analysis compared to the single-study discovery EWAS (Section 2.5). Finally, we did not conduct orthogonal or wet-lab validation of replicated sex DMPs. Such steps would be essential to narrow biomarker candidates and elucidate biological mechanisms explaining differential DNAm.

Our cross-study and cross-platform approach could be applied to other highly prevalent tissues such as brain [1] or expanded to include public bisulfite-sequencing samples from the Sequence Read Archive [65], which could clarify the genome region specificity of high-confidence biomarkers from DNAm arrays [7, 66, 67]. A future update of `recountmethylation` may include such samples.

## 4 Methods

### 4.1 Compiling recent public DNAm array data across platforms

DNAm array data were identified, downloaded, and processed using the `recountmethylation_instance` v0.0.1 [54] Snakemake workflow. It comprises the `recountmethylation_server` v1.0.0 [52] and `recountmethylation.pipeline` v1.0.0 [53] tools, which we used previously to compile HM450K data [1]. We uniformly processed samples into cross-study and cross-platform data compilations. Data were from samples run on the HM450K [4] and EPIC [7] DNAm array platforms and available on GEO by March 31, 2021. Sample DNAm was normalized using out of band signal (a.k.a. “noob-normalization” [49]). Compilations paired normalized DNAm fractions, or Beta-values, with harmonized sample metadata for 68,758 cumulative samples for which raw image data files, or IDATs, were available as gzip-compressed supplementary files.

Compilations were stored as Hierarchical Data Format 5 (HDF5)-based Summarized-Experiment files generated using the `HDF5Array` v1.18.0 and `rhdf5` v2.34.0 R/Bioconductor packages [68, 69]. These formats used `DelayedArray` v0.18.0 [70] to support rapid access, summaries, and filters. For most analyses, DNAm data were merged across platforms for the 453,093 CpG probes [7] they shared. We made compiled data available online at https://recount.bio/data/gr-gseadj_h5se_hm450k-epic-merge_0-0-3/.

### 4.2 Prediction and harmonization of sample metadata

We generated harmonized sample metadata from heterogeneous metadata mined from SOFT files accompanying GEO studies. We wrote regex terms to detect keywords in metadata-containing files, and we mapped detected terms to controlled vocabularies under “tissue”, “disease,” and other categories, as described in [1] and [71]. The suitability of regex patterns for capturing informative metadata terms was spot-checked across studies and iteratively updated to more precisely map terms and avoid erroneous mappings. We further predicted sample types from mined metadata using the method from [72]. To add sex annotations, we used the `minfi` v1.37.1 [50, 73]) R package; to add six blood type cell fractions, we used the method from [38]); and to add age annotations, we used the pan-tissue epigenetic clock model from [26]. Finally, we calculated the top components of genetic ancestry using the method from [41].

### 4.3 Sample QC filters

We used metadata filters to find the 3 most prevalent blood sample types (whole blood, cord blood, and PBMCs), and the aggregate type “all,” which includes the above types and samples of whose type was indeterminate from mined metadata. We then performed QC with reference to prior findings from [1]. We removed samples for which either: (1) log2 median M and U signals were both *<*10; or (2) the sample failed *≥*2/5 most informative BeadArray metrics. These criteria removed 245 samples, and all but one was run on the HM450K platform.

### 4.4 Simulation of study bias adjustments

We used simulations to show the impact of study ID adjustment on explained variance. As detailed in Fig. S1, simulations consisted of 4 steps: (1) calculate sample DNAm M-values from 500 CpG probes and 5 studies, selected randomly; (2) adjust study ID across all 5 selected studies (i.e. “adjustment 1”) or subsets of 2-4 studies (i.e. “adjustment 2”); (3) perform ANOVA for 3 models; (4) get FEV for each variable across 3 models. In total, simulations used 29,028 unique CpG probes and 62 unique studies.

Multiple regression models accounted for sample type, platform, study ID, DNAm-based predictions for age, sex, and six cell type fractions, and two genetic ancestry components, which were determined as described above. Variables were grouped as one of biological (i.e. six blood cell type fractions), demographic (i.e. age, sex, and two genetic ancestry components), and technical (i.e. platform). Study bias adjustments were performed using the `removeBatchEffect()` function from the `limma` v3.46.0 [74] R package. Parallel sessions were deployed using the `parallel` v4.1.1 R package.

### 4.5 PCA of autosomal DNAm

We performed autosomal DNAm PCA on compiled blood samples using a reduced 1,000-dimensional representation of the normalized and bias-corrected Beta-values [75, 76] obtained via feature hashing. (See [1] for details on this approach.) For the top 10 components, we calculated FEV from ANOVA using multiple regression models containing the 13 variables from the 3 categories described above.

### 4.6 Blood autosomal DNAm search index construction

We used the `hnswlib` v0.5.2 Python library to make a DNAm-based search index from which one can rapidly identify the nearest samples which neighbor one or more queried DNAm profiles [55]. We used the Hierarchical and Navigable Small Worlds (HNSW) graph algorithm implemented in `hnswlib`, as this was among the top performing algorithms from a recent comprehensive benchmark of search algorithms [77]. With the `mmh3` v3.0.0 and `numpy` v1.20.1 [78] Python libraries, we applied feature hashing to generate a reduced 1,000-dimensional representation of each sample [76, 79] of each blood sample’s noob-normalized Beta-values. The search index files are available online at https://recount.bio/data/sindexhnsw_bval-gseadj-fh10k_all-blood-2-platforms.pickle and https://recount.bio/data/sidicthnsw_bval-gseadj-fh10k_all-blood-2-platforms.pickle.

### 4.7 Power analyses using pwrEWAS

We used the method provided in the `pwrEWAS` v1.4.0 R/Bioconductor library to perform power analyses across DNAm array platforms [42]. Parameters for these analyses included 100 total simulations varying the total samples *N* from 50 to 850. We targeted 500 DMPs and assessed test group Beta-value differences *δ* of 0.05, 0.1, and 0.2.

### 4.8 Replication of whole blood sex DMPs

We replicated sex DMPs from [48], a study of whole blood from Japanese individuals, using independent compilations of whole blood and PBMC samples in `recountmethylation`. After filtering out sex chromosome and cross-reactive probes [80], there were 375,244 CpG probes in whole blood and 375,244 CpG probes in PBMCs. After filtering for sample quality, we used data from 5,980 whole blood samples (3,942 females and 2,924 males) and 642 PBMC samples (397 females and 230 males). Ages tended towards young adult and middle-aged for whole blood (age, mean*±*SD, 39*±*21 years) and samples from [48] (46*±*12 years), but were more frequently from adolescents and young adults among PBMC (25*±*19 years). We preprocessed DNAm M-values using surrogate variables analysis (SVA) with the `sva` v3.4.0 R package [81]. We determined sex DMPs using coefficient P-values for the sex variable in multiple regressions, where regression models used biological, demographic, and technical variables described above (Supplementary Materials).

### 4.9 Statistical analyses and visualizations

Data processing and analyses were performed using the R v4.1.0 and Python v3.7.1 programming languages [82, 83]. Statistical summaries and tests were performed using base R libraries. DNAm array processing, normalization, analysis, and prediction of sex and six blood cell type fractions was performed using the `minfi`, `minfiData`, and `minfiDataEPIC` R packages. Workflow diagrams were created using BioRender.com. Visualizations in Results made us of the `ggplot2` v3.3.2, `grid` v4.1.3, `gridExtra` v2.3, `UpSetR` v1.4.0, `ggpubr` v0.4.0, `ggforce` v0.3.3, and `png` v0.1-7 R packages [84, 85]. P-value adjustments used either the Bonferroni method or the Benjamini-Hotchberg method [86]. Enrichment tests used the `binom.test()` base R function with the background of 453,093 total probes overlapping both array platforms [82]. Supplemental scripts and functions recreating our results are available online at https://www.github.com/metamaden/recountmethylation_v2_manuscript.

### 4.10 Supplemental data, files, and code

The following resources have been provided to reproduce results, figures, and tables in this paper:

1. The updated `recountmethylation` Bioconductor package is now available (https://doi.org/doi:10.18129/B9.bioc.recountmethylation). It features new functions supporting analysis of large data compilations, and new vignettes showing how to perform novel power analysis, infer genetic ancestry, and more using DNAm array data.
2. Supplemental code and scripts for this paper, including support for creating and querying a search index of DNAm array samples, are available in the manuscript GitHub repository (https://github.com/metamaden/recountmethylation_v2_manuscript).
3. The `recountmethylation instance` Snakemake workflow is available on GitHub [54]. This will be useful for researchers hoping to make and update new compilations of public DNAm array data from GEO.

## Acknowledgments

We thank Joe Gray, Jeremy Goecks, and Stephanie Hicks for early feedback on the content of this manuscript.

## Disclaimer

The contents do not represent the views of the U.S. Department of Veterans Affairs or the United States Government.

## Supplementary information

### Supplementary figures

**Fig. S1:**
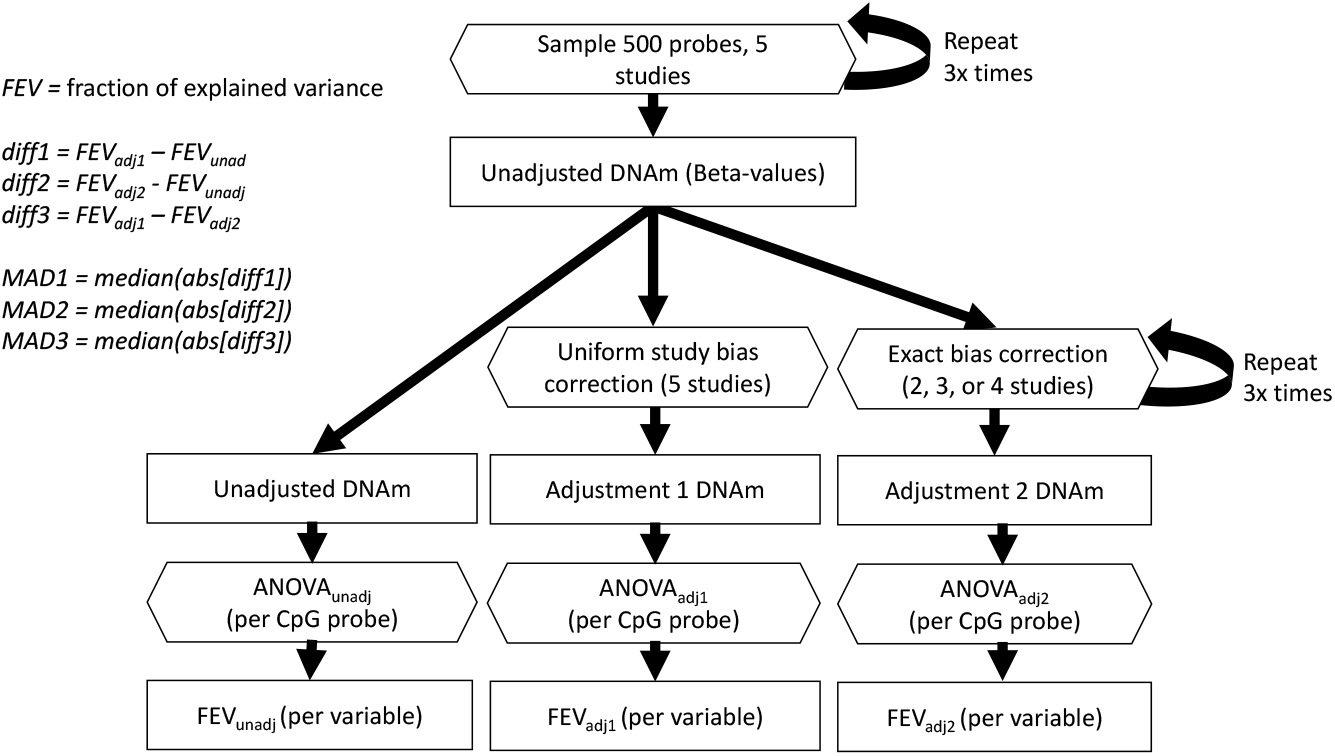
Workflow diagram to simulate the impact of GSE bias corrections on explained variances. This diagram shows a single simulation rep, including repeated probe and study selections where indicated. From top to bottom, the workflow shows random selection of 500 CpG probes, random selection of 5 studies, and calculation of 3 DNAm datasets per CpG probe: (1) unadjusted DNAm; (2) DNAm after local adjustment on subsets between 1-4 study IDs among 5 selected (a.k.a. adjustment 1); (3) DNAm after uniform adjustment on all 5 selected study IDs (a.k.a. adjustment 2). Finally, ANOVAs are conducted across the 3 DNAm models, and fractions of explained variances are determined from sum of squared variances (Methods). Terminology for workflow terms is shown at top left.

**Fig. S2:**
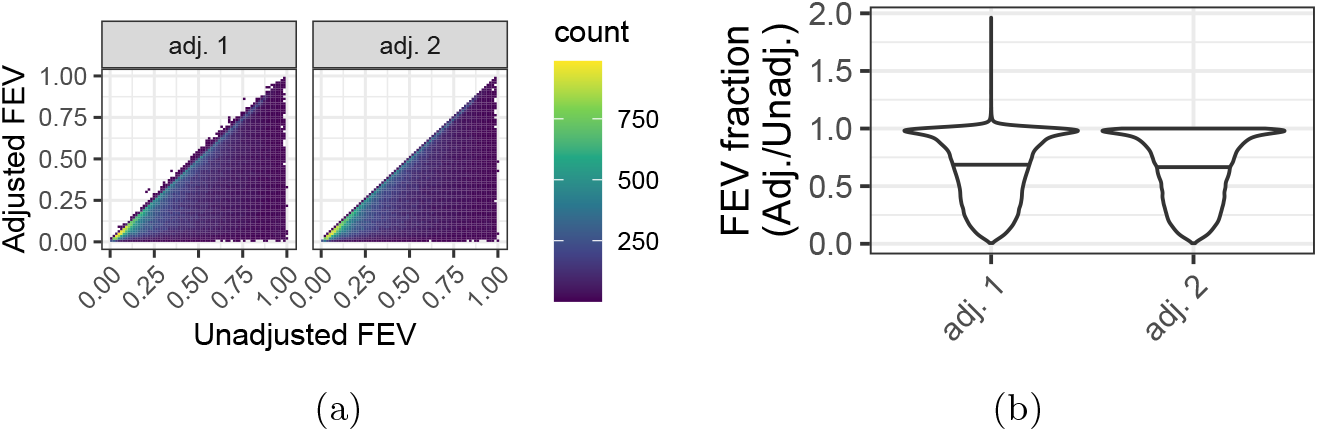
Fraction of explained variance (FEV) between unadjusted and adjusted DNAm across GSE bias simulations. (a) Density plots of unadjusted FEV on x axes and adjusted FEV on y-axes. Color fill shows density of simulation outcome counts (dark blue = low, green = moderate, yellow = high). (b) Violin plots of FEV fractions, or adjusted FEV over unadjusted FEV, by adjustment type on the x-axis.

**Fig. S3:**
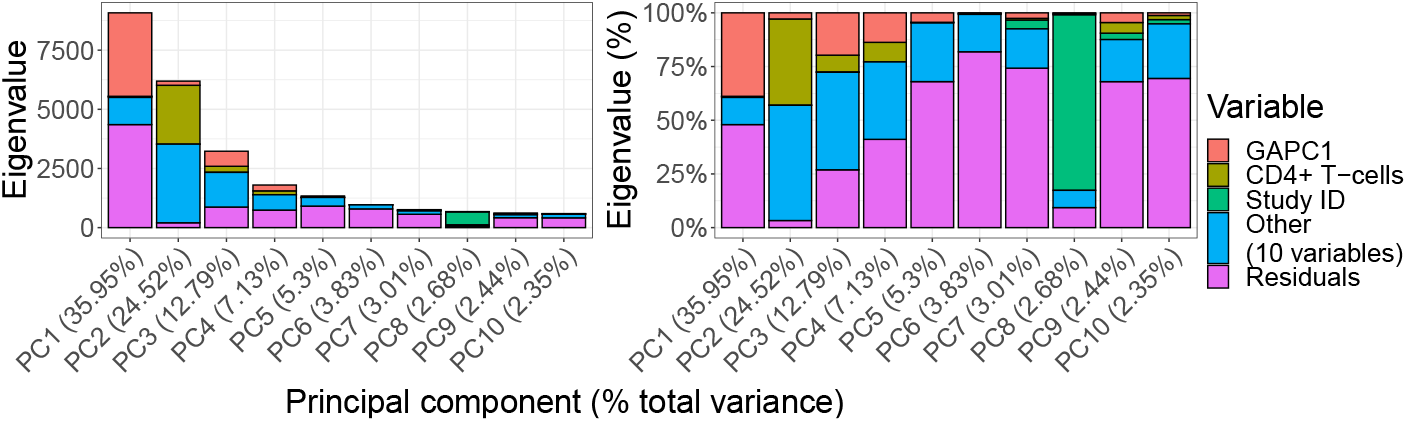
Autosomal DNAm PCA results across normal blood samples. Eigenvalues explained by select variables. Stacked barplots show the eigenvalue magnitudes (left) and percentages (right) for the top ten components (x-axis). Fill colors indicate magnitudes of component sum of squared variances explained by select variables (red = genetic ancestry PC1, yellow = predicted CD4+ T-cells, green = Study ID, blue = other variable, purple = residuals). The term “other” stores the 10 remaining model variables tested. X-axis labels show the percent of total variances explained by each component in parentheses.

**Fig. S4:**
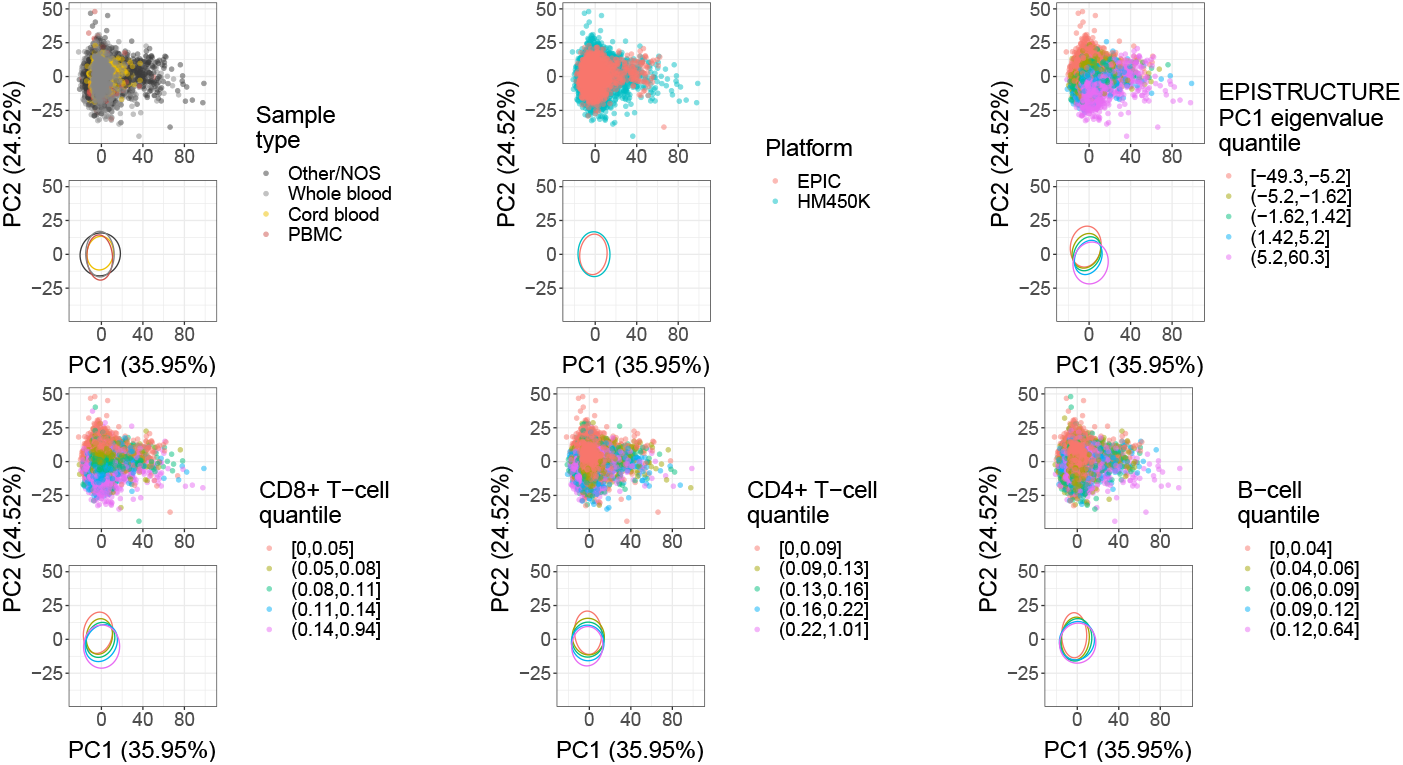
Top two components from PCAs of autosomal DNAm (noob-normalized Beta-values) colored according to different variables. Each panel includes a scatter plot (top) and 95% confidence interval ellipses (bottom) where the x and y axes correspond to, respectively, the first and second components. Colors specify sample type (top left, black = other/not otherwise specified, gray = whole blood, yellow = cord blood, red = PBMC), platform (top middle, red = EPIC, blue = HM450K), the first component of genetic ancestry (top right, [41]), and predicted fractions [38] for CD8+ T-cells (bottom left), CD4+ T-cells (bottom middle), and B-cells (bottom right). Color labels for the latter four continuous variables correspond to sample quintile bins (e.g. 5 quantile ranges: pink = 0-20, yellow = 20-40, green = 40-60, blue = 60-80, purple = 80-100).

**Fig. S5:**
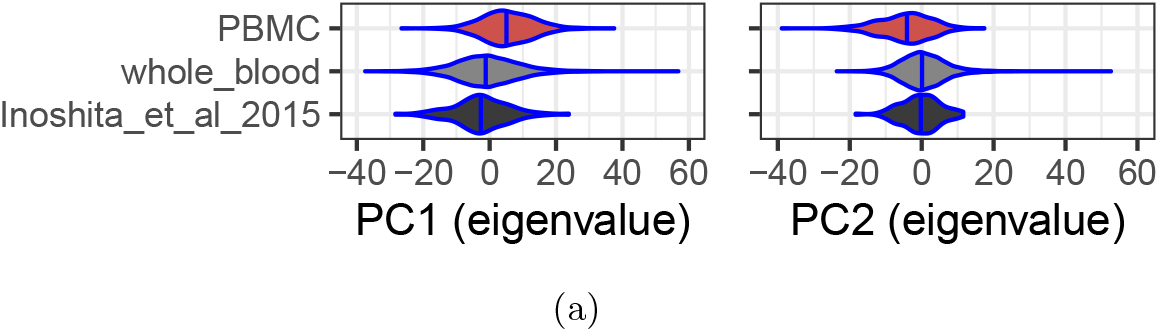
Violin plots of the top two genetic ancestry components on the x axes for each of the three available datasets on the y axes (black = “Inoshita et al 2015” [48], red = PBMC compilation, gray = whole blood compilation). Vertical blue lines represent the distribution medians.

**Fig. S6:**
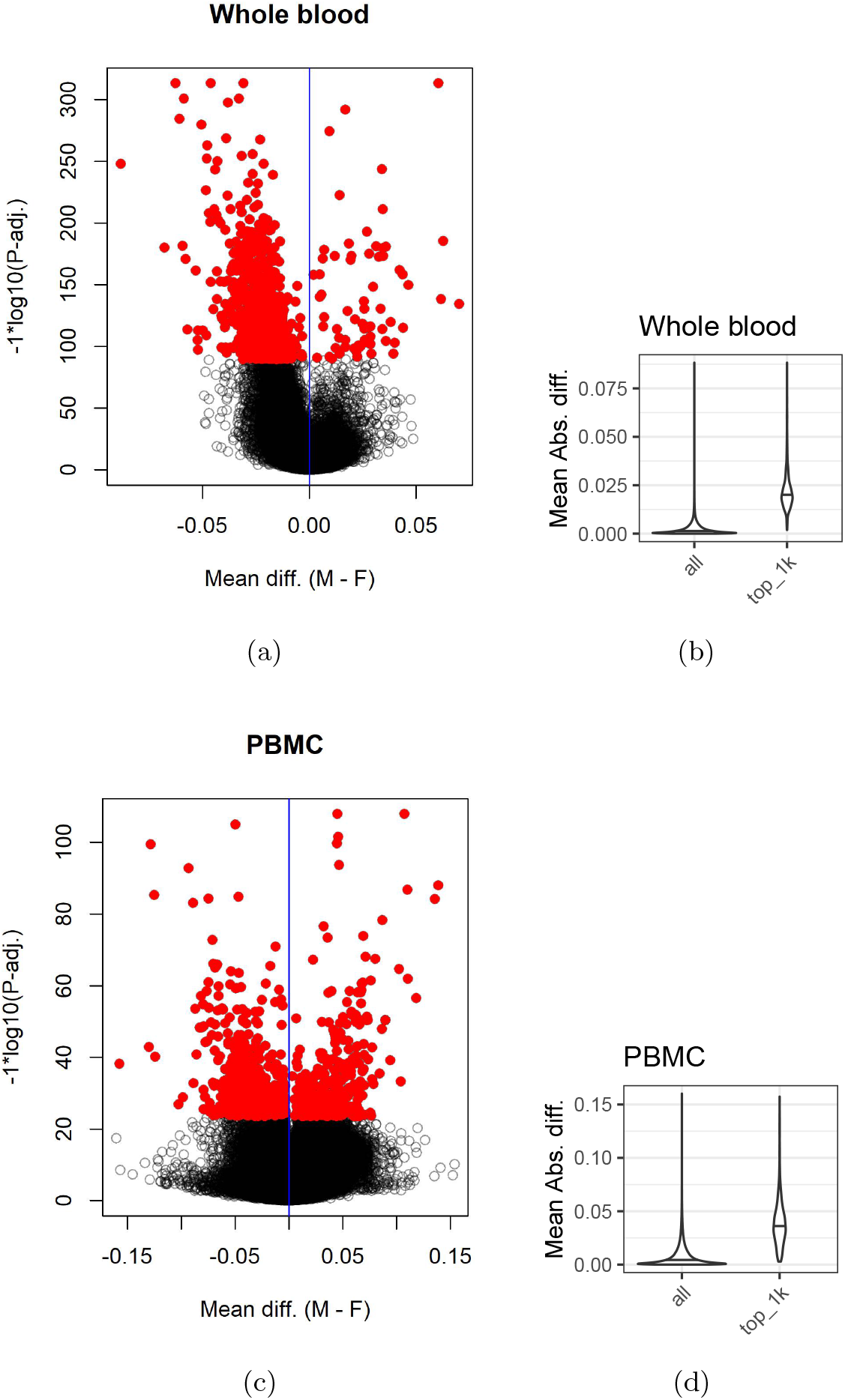
Summaries of differential DNAm by sex in whole blood and peripheral blood mononuclear cells (PBMC). (a) Volcano plot for whole blood showing difference in mean Beta-values (males minus females, x-axis) versus probe significance (−1*log10[P-adj.], Benjamini Hotchberg adjustment [86], y-axis). Red dots represent the top 1,000 DMPs, black circles represent non-DMP probes. (b) violin plots for whole blood showing distributions of absolute difference in mean Beta-values (males minus females, y-axis) for all tested CpG probes (left) and only the top 1,000 most significant differentially methylated probes (DMPs, right). Horizontal black lines indicate the distribution medians. (c) Volcano plot for PBMC showing difference in mean Beta-values (males minus females, x-axis) versus probe significance (−1*log10[P-adj.], y-axis). Red dots represent the top 1,000 DMPs, black circles represent non-DMP probes. (d) violin plots for PBMC showing distributions of absolute difference in mean Beta-values (males minus females, y-axis) for all tested CpG probes (left) and only the top 1,000 most significant differentially methylated probes (DMPs, right). Horizontal black lines indicate the distribution medians.

**Fig. S7:**
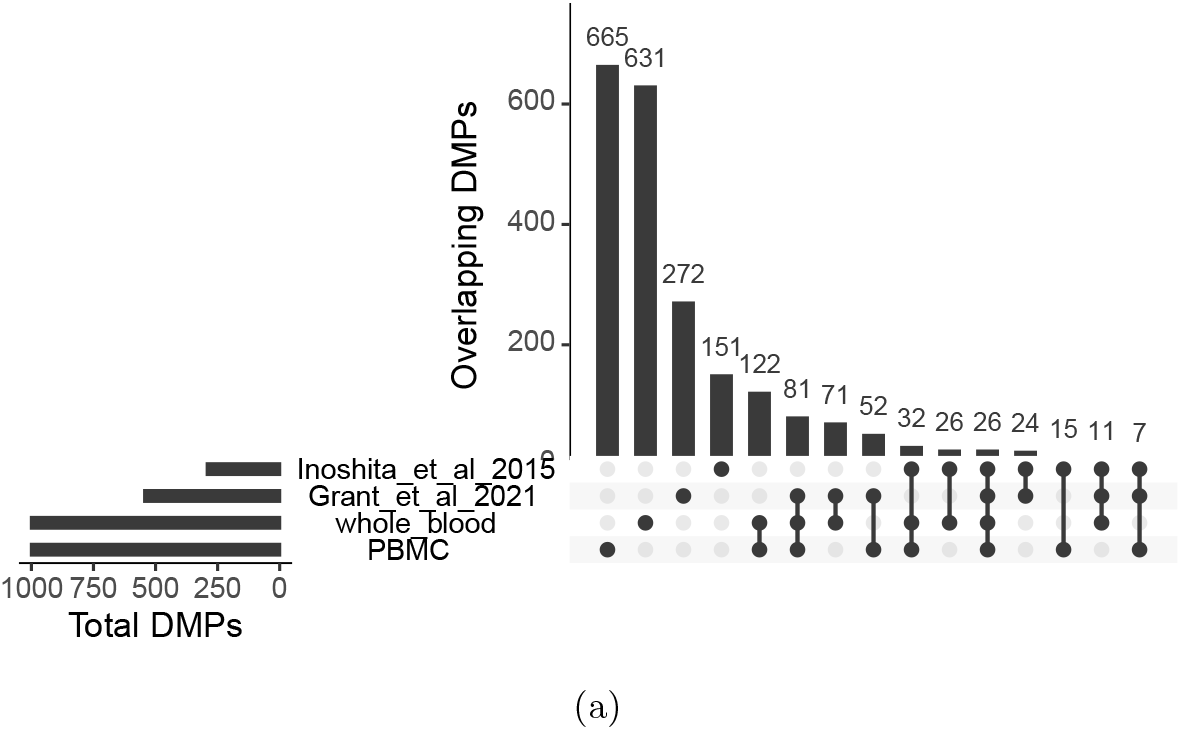
Upset plot showing DMP overlaps (lower left and top right barplot magnitudes) among 4 DMP sets including compiled whole blood, compiled PBMC, “Inoshita et al 2015” [48], and “Grant et al 2021” [47] (lower y-axis labels). Set magnitudes are shown on top of subset magnitude barplots.

### Supplementary tables

**Table S1:**
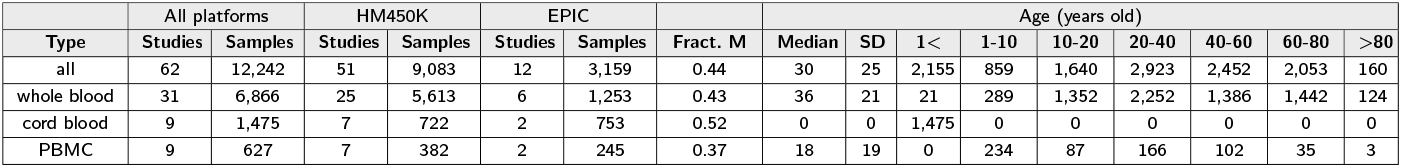
Blood sample type (rows) availability and demographic variables (columns).

**Table S2:** Fraction of explained variance (FEV) medians by model type (rows) and variables (columns), across study ID bias correction simulations (Methods).

| Model | Technical |  | Biological |  |  |  |  |  | Demographic |  |  |  |
| --- | --- | --- | --- | --- | --- | --- | --- | --- | --- | --- | --- | --- |
|  | Study ID | Platform | CD8+ T-cells | CD4+ T-cells | B-cells | Gran. | Mono. | Natural Killer | Sex | Age | G.A. PC1 | G.A. PC2 |
| Unadj. | 0.465 | 0.037 | 0.014 | 0.010 | 0.013 | 0.020 | 0.005 | 0.009 | 0.003 | 0.012 | 0.077 | 0.054 |
| Adj. 1 | 0.002 | 0.086 | 0.036 | 0.026 | 0.035 | 0.050 | 0.013 | 0.024 | 0.008 | 0.030 | 0.212 | 0.144 |
| Adj. 2 | 0.000 | 0.089 | 0.037 | 0.026 | 0.035 | 0.051 | 0.013 | 0.024 | 0.008 | 0.031 | 0.214 | 0.144 |

## References

[1] Maden, S. K., Thompson, R. F., Hansen, K. D. & Nellore, A. Human methylome variation across Infinium 450K data on the Gene Expression Omnibus. NAR Genomics and Bioinformatics 3 (2) (2021). URL https://doi.org/10.1093/nargab/lqab025. https://doi.org/10.1093/nargab/lqab025.

[2] Edgar, R., Domrachev, M. & Lash, A. E. Gene Expression Omnibus: NCBI gene expression and hybridization array data repository. Nucleic Acids Research 30 (1), 207–210 (2002). URL https://doi.org/10.1093/nar/30.1.207. https://doi.org/10.1093/nar/30.1.207.

[3] Barrett, T. et al. Ncbi geo: archive for functional genomics data sets—update. Nucleic acids research 41 (D1), D991–D995 (2012).

[4] Bibikova, M. et al. High density DNA methylation array with single CpG site resolution. Genomics 98 (4), 288–295 (2011). URL http://www.sciencedirect.com/science/article/pii/S0888754311001807. https://doi.org/10.1016/j.ygeno.2011.07.007.

[5] Sandoval, J. et al. Validation of a DNA methylation microarray for 450,000 CpG sites in the human genome. Epigenetics 6 (6), 692–702 (2011). https://doi.org/10.4161/epi.6.6.16196.

[6] Huber, W. et al. Orchestrating high-throughput genomic analysis with Bioconductor. Nature Methods 12 (2), 115–121 (2015). URL https://www.nature.com/articles/nmeth.3252. https://doi.org/10.1038/nmeth.3252, number: 2 Publisher: Nature Publishing Group.

[7] Pidsley, R. et al. Critical evaluation of the Illumina MethylationEPIC BeadChip microarray for whole-genome DNA methylation profiling. Genome Biology 17 (2016). URL https://www.ncbi.nlm.nih.gov/pmc/articles/PMC5055731/. https://doi.org/10.1186/s13059-016-1066-1.

[8] Li, L. et al. DNA methylation in peripheral blood: a potential biomarker for cancer molecular epidemiology. Journal of Epidemiology 22 (5), 384–394 (2012). https://doi.org/10.2188/jea.je20120003.

[9] Locke, W. J. et al. DNA Methylation Cancer Biomarkers: Translation to the Clinic. Frontiers in Genetics 10, 1150 (2019). URL https://www.frontiersin.org/article/10.3389/fgene.2019.01150. https://doi.org/10.3389/fgene.2019.01150.

[10] Mikeska, T. & Craig, J. M. DNA Methylation Biomarkers: Cancer and Beyond. Genes 5 (3), 821–864 (2014). URL https://www.ncbi.nlm.nih.gov/pmc/articles/PMC4198933/. https://doi.org/10.3390/genes5030821.

[11] Willmer, T., Johnson, R., Louw, J. & Pheiffer, C. Blood-Based DNA Methylation Biomarkers for Type 2 Diabetes: Potential for Clinical Applications. Frontiers in Endocrinology 9, 744 (2018). URL https://www.frontiersin.org/article/10.3389/fendo.2018.00744. https://doi.org/10.3389/fendo.2018.00744.

[12] Bacos, K. et al. Blood-based biomarkers of age-associated epigenetic changes in human islets associate with insulin secretion and diabetes. Nature Communications 7, 11089 (2016). URL https://www.ncbi.nlm.nih.gov/pmc/articles/PMC4821875/. https://doi.org/10.1038/ncomms11089.

[13] Dayeh, T. et al. DNA methylation of loci within ABCG1 and PHOSPHO1 in blood DNA is associated with future type 2 diabetes risk. Epigenetics 11 (7), 482–488 (2016). URL https://www.ncbi.nlm.nih.gov/pmc/articles/PMC4939923/. https://doi.org/10.1080/15592294.2016.1178418.

[14] Samblas, M., Milagro, F. I. & Martinez, A. DNA methylation markers in obesity, metabolic syndrome, and weight loss. Epigenetics 14 (5), 421–444 (2019). URL https://www.ncbi.nlm.nih.gov/pmc/articles/PMC6557553/. https://doi.org/10.1080/15592294.2019.1595297.

[15] Hyun, J. & Jung, Y. DNA Methylation in Nonalcoholic Fatty Liver Disease. International Journal of Molecular Sciences 21 (21), 8138 (2020). URL https://www.ncbi.nlm.nih.gov/pmc/articles/PMC7662478/. https://doi.org/10.3390/ijms21218138.

[16] Hudon Thibeault, A.-A. & Laprise, C. Cell-Specific DNA Methylation Signatures in Asthma. Genes 10 (11), 932 (2019). URL https://www.ncbi.nlm.nih.gov/pmc/articles/PMC6896152/. https://doi.org/10.3390/genes10110932.

[17] Fransquet, P. D. et al. Blood DNA methylation signatures to detect dementia prior to overt clinical symptoms. Alzheimer’s & Dementia: Diagnosis, Assessment & Disease Monitoring 12 (1), e12056 (2020). URL https://onlinelibrary.wiley.com/doi/abs/10.1002/dad2.12056. https://doi.org/10.1002/dad2.12056, eprint: https://onlinelibrary.wiley.com/doi/pdf/10.1002/dad2.12056.

[18] Jensen, S. O. et al. Novel DNA methylation biomarkers show high sensitivity and specificity for blood-based detection of colorectal cancer—a clinical biomarker discovery and validation study. Clinical Epigenetics 11 (1), 158 (2019). URL https://doi.org/10.1186/s13148-019-0757-3. https://doi.org/10.1186/s13148-019-0757-3.

[19] Dong, L. & Ren, H. Blood-based DNA Methylation Biomarkers for Early Detection of Colorectal Cancer. Journal of proteomics & bioinformatics 11 (6), 120–126 (2018). URL https://www.ncbi.nlm.nih.gov/pmc/articles/PMC6054487/. https://doi.org/10.4172/jpb.1000477.

[20] Alizadeh-Sedigh, M., Fazeli, M. S., Mahmoodzadeh, H., Sharif, S. B. & Teimoori-Toolabi, L. Methylation of FBN1, SPG20, ITF2, RUNX3, SNCA, MLH1, and SEPT9 genes in circulating cell-free DNA as biomarkers of colorectal cancer. Cancer Biomarkers: Section A of Disease Markers (2021). https://doi.org/10.3233/CBM-210315

[21] Lin, W.-H. et al. Circulating tumor DNA methylation marker MYO1-G for diagnosis and monitoring of colorectal cancer. Clinical Epigenetics 13 (1), 232 (2021). https://doi.org/10.1186/s13148-021-01216-0.

[22] Yu, M. et al. Subtypes of Barrett’s oesophagus and oesophageal adenocarcinoma based on genome-wide methylation analysis. Gut (2018). https://doi.org/10.1136/gutjnl-2017-314544.

[23] Guan, Z., Yu, H., Cuk, K., Zhang, Y. & Brenner, H. Whole-Blood DNA Methylation Markers in Early Detection of Breast Cancer: A Systematic Literature Review. Cancer Epidemiology and Prevention Biomarkers 28 (3), 496–505 (2019). URL https://cebp.aacrjournals.org/content/28/3/496. https://doi.org/10.1158/1055-9965.EPI-18-0378, publisher: American Association for Cancer Research Section: Minireview.

[24] Henriksen, S. D. & Thorlacius-Ussing, O. Cell-Free DNA Methylation as Blood-Based Biomarkers for Pancreatic Adenocarcinoma—A Literature Update. Epigenomes 5 (2), 8 (2021). URL https://www.mdpi.com/2075-4655/5/2/8. https://doi.org/10.3390/epigenomes5020008, number: 2 Publisher: Multidisciplinary Digital Publishing Institute.

[25] Danstrup, C. S. et al. DNA methylation biomarkers in peripheral blood of patients with head and neck squamous cell carcinomas. A systematic review. PLOS ONE 15 (12), e0244101 (2020). URL https://journals.plos.org/plosone/article?id=10.1371/journal.pone.0244101. https://doi.org/10.1371/journal.pone.0244101, publisher: Public Library of Science.

[26] Horvath, S. DNA methylation age of human tissues and cell types. Genome Biology 14 (10), R115 (2013). URL http://www.ncbi.nlm.nih.gov/pmc/articles/PMC4015143/. https://doi.org/10.1186/gb-2013-14-10-r115.

[27] Hannum, G. et al. Genome-wide Methylation Profiles Reveal Quantitative Views of Human Aging Rates. Molecular Cell 49 (2), 359–367 (2013). URL https://www.cell.com/molecular-cell/abstract/S1097-2765(12)00893-3. https://doi.org/10.1016/j.molcel.2012.10.016.

[28] Merid, S. K. et al. Epigenome-wide meta-analysis of blood DNA methylation in newborns and children identifies numerous loci related to gestational age. Genome Medicine 12 (2020). URL https://www.ncbi.nlm.nih.gov/pmc/articles/PMC7050134/. https://doi.org/10.1186/s13073-020-0716-9.

[29] Haftorn, K. L. et al. An EPIC predictor of gestational age and its application to newborns conceived by assisted reproductive technologies. Clinical Epigenetics 13 (1), 82 (2021). URL https://doi.org/10.1186/s13148-021-01055-z. https://doi.org/10.1186/s13148-021-01055-z.

[30] Huang, Y.-T. et al. Epigenome-wide profiling of DNA methylation in paired samples of adipose tissue and blood. Epigenetics 11 (3), 227–236 (2016). URL https://doi.org/10.1080/15592294.2016.1146853. https://doi.org/10.1080/15592294.2016.1146853, publisher: Taylor & Francis eprint: https://doi.org/10.1080/15592294.2016.1146853.

[31] Åsenius, F. et al. The DNA methylome of human sperm is distinct from blood with little evidence for tissue-consistent obesity associations. PLOS Genetics 16 (10), e1009035 (2020). URL https://journals.plos.org/plosgenetics/article?id=10.1371/journal.pgen.1009035. https://doi.org/10.1371/journal.pgen.1009035, publisher: Public Library of Science.

[32] Parveen, N. & Dhawan, S. DNA Methylation Patterning and the Regulation of Beta Cell Homeostasis. Frontiers in Endocrinology 12, 651258 (2021). URL https://www.ncbi.nlm.nih.gov/pmc/articles/PMC8137853/. https://doi.org/10.3389/fendo.2021.651258.

[33] Mayne, B. T. et al. Accelerated placental aging in early onset preeclampsia pregnancies identified by DNA methylation. Epigenomics 9 (3), 279–289 (2017). URL https://www.futuremedicine.com/doi/full/10.2217/epi-2016-0103. https://doi.org/10.2217/epi-2016-0103, publisher: Future Medicine.

[34] Knight, A. K. et al. An epigenetic clock for gestational age at birth based on blood methylation data. Genome Biology 17 (1), 206 (2016). URL https://doi.org/10.1186/s13059-016-1068-z. https://doi.org/10.1186/s13059-016-1068-z.

[35] Bohlin, J. et al. Prediction of gestational age based on genome-wide differentially methylated regions. Genome Biology 17 (1), 207 (2016). URL https://doi.org/10.1186/s13059-016-1063-4. https://doi.org/10.1186/s13059-016-1063-4.

[36] Lee, Y. et al. Placental epigenetic clocks: estimating gestational age using placental DNA methylation levels. Aging (Albany NY) 11 (12), 4238–4253 (2019). URL https://www.ncbi.nlm.nih.gov/pmc/articles/PMC6628997/. https://doi.org/10.18632/aging.102049.

[37] Fung, R. et al. Achieving accurate estimates of fetal gestational age and personalised predictions of fetal growth based on data from an international prospective cohort study: a population-based machine learning study. The Lancet. Digital Health 2 (7), e368–e375 (2020). URL https://www.ncbi.nlm.nih.gov/pmc/articles/PMC7323599/. https://doi.org/10.1016/S2589-7500(20)30131-X.

[38] Houseman, E. A. et al. DNA methylation arrays as surrogate measures of cell mixture distribution. BMC Bioinformatics 13 (1), 86 (2012). URL http://www.biomedcentral.com/1471-2105/13/86/abstract. https://doi.org/10.1186/1471-2105-13-86.

[39] Koestler, D. C. et al. Improving cell mixture deconvolution by identifying optimal DNA methylation libraries (IDOL). BMC Bioinformatics 17 (2016). URL https://www.ncbi.nlm.nih.gov/pmc/articles/PMC4782368/. https://doi.org/10.1186/s12859-016-0943-7

[40] Salas, L. A. et al. Enhanced cell deconvolution of peripheral blood using DNA methylation for high-resolution immune profiling. Nature Communications 13 (1), 761 (2022). URL https://www.nature.com/articles/s41467-021-27864-7. https://doi.org/10.1038/s41467-021-27864-7, number: 1 Publisher: Nature Publishing Group

[41] Rahmani, E. et al. Genome-wide methylation data mirror ancestry information. Epigenetics & Chromatin 10 (1), 1 (2017). URL https://doi.org/10.1186/s13072-016-0108-y. https://doi.org/10.1186/s13072-016-0108-y.

[42] Graw, S., Henn, R., Thompson, J. A. & Koestler, D. C. pwrEWAS: a user-friendly tool for comprehensive power estimation for epigenome wide association studies (EWAS). BMC Bioinformatics 20 (1), 218 (2019). URL https://doi.org/10.1186/s12859-019-2804-7. https://doi.org/10.1186/s12859-019-2804-7.

[43] Masser, D. R. et al. Sexually divergent DNA methylation patterns with hippocampal aging. Aging Cell 16 (6), 1342–1352 (2017). URL https://www.ncbi.nlm.nih.gov/pmc/articles/PMC5676057/. https://doi.org/10.1111/acel.12681.

[44] Hall, E. et al. Sex differences in the genome-wide DNA methylation pattern and impact on gene expression, microRNA levels and insulin secretion in human pancreatic islets. Genome Biology 15 (12), 522 (2014). URL https://doi.org/10.1186/s13059-014-0522-z. https://doi.org/10.1186/s13059-014-0522-z.

[45] Nino, C. L. et al. Characterization of Sex-Based Dna Methylation Signatures in the Airways During Early Life. Scientific Reports 8, 5526 (2018). URL https://www.ncbi.nlm.nih.gov/pmc/articles/PMC5882800/. https://doi.org/10.1038/s41598-018-23063-5

[46] Maschietto, M. et al. Sex differences in DNA methylation of the cord blood are related to sex-bias psychiatric diseases. Scientific Reports 7 (1), 44547 (2017). URL https://www.nature.com/articles/srep44547. https://doi.org/10.1038/srep44547, bandiera abtest: a Cc license type: cc by Cg type: Nature Research Journals Number: 1 Primary atype: Research Publisher: Nature Publishing Group Subject term: DNA methylation;Stress and resilience Subject term id: dna-methylation;stress-and-resilience.

[47] Grant, O. A., Wang, Y., Kumari, M., Zabet, N. R. & Schalkwyk, L. Characterising sex differences of autosomal DNA methylation in whole blood using the Illumina EPIC array. bioRxiv 2021.09.02.458717 (2021). https://doi.org/10.1101/2021.09.02.458717, company: Cold Spring Harbor Laboratory Distributor: Cold Spring Harbor Laboratory Label: Cold Spring Harbor Laboratory Section: New Results Type: article.

[48] Inoshita, M. et al. Sex differences of leukocytes DNA methylation adjusted for estimated cellular proportions. Biology of Sex Differences 6 (1), 11 (2015). URL https://doi.org/10.1186/s13293-015-0029-7. https://doi.org/10.1186/s13293-015-0029-7.

[49] Triche, T. J., Weisenberger, D. J., Van Den Berg, D., Laird, P. W. & Siegmund, K. D. Low-level processing of Illumina Infinium DNA Methylation BeadArrays. Nucleic Acids Research 41 (7), e90–e90 (2013). URL https://academic.oup.com/nar/article/41/7/e90/1070878. https://doi.org/10.1093/nar/gkt090.

[50] Aryee, M. J. et al. Minfi: A flexible and comprehensive bioconductor package for the analysis of infinium dna methylation microarrays. Bioinformatics 30 (10), 1363–1369 (2014). https://doi.org/10.1093/bioinformatics/btu049.

[51] Murray, J. R. & Rajeevan, M. S. Evaluation of DNA extraction from granulocytes discarded in the separation medium after isolation of peripheral blood mononuclear cells and plasma from whole blood. BMC Research Notes 6, 440 (2013). URL https://www.ncbi.nlm.nih.gov/pmc/articles/PMC3818442/. https://doi.org/10.1186/1756-0500-6-440.

[52] Maden, S., Thompson, R., Hansen, K. & Nellore, A. recountmethylation server (2021). URL https://github.com/metamaden/recountmethylation server. Recountmethylation server Python package for DNAm array queries and downloads from GEO.

[53] Maden, S., Thompson, R., Hansen, K. & Nellore, A. recountmethylation.pipeline (2021). URL https://github.com/metamaden/recountmethylation.pipeline. Recountmethylation.pipeline R package for uniformly processing and harmonoizing cross-study compilations of DNAm array data.

[54] Maden, S., Thompson, R., Hansen, K. & Nellore, A. recountmethylation instance (2022). URL https://github.com/metamaden/recountmethylation instance. Recountmethylation instance Snakemake workflow for synchronization of DNAm array data from GEO.

[55] Malkov, Y. A. & Yashunin, D. A. Efficient and robust approximate nearest neighbor search using Hierarchical Navigable Small World graphs. 1603.09320 [cs] (2018). URL http://arxiv.org/abs/1603.09320. ArXiv: 1603.09320.

[56] Mölder, F. et al. Sustainable data analysis with Snakemake. F1000Research 10 (10:33) (2021). URL https://f1000research.com/articles/10-33. https://doi.org/10.12688/f1000research.29032.1.

[57] Mansell, G. et al. Guidance for DNA methylation studies: statistical insights from the Illumina EPIC array. BMC Genomics 20 (2019). URL https://www.ncbi.nlm.nih.gov/pmc/articles/PMC6518823/. https://doi.org/10.1186/s12864-019-5761-7.

[58] Gardiner-Garden, M. & Frommer, M. CpG islands in vertebrate genomes. Journal of Molecular Biology 196 (2), 261–282 (1987). https://doi.org/10.1016/0022-2836(87)90689-9.

[59] Takai, D. & Jones, P. A. Comprehensive analysis of CpG islands in human chromosomes 21 and 22. Proceedings of the National Academy of Sciences of the United States of America 99 (6), 3740–3745 (2002). https://doi.org/10.1073/pnas.052410099.

[60] Deaton, A. M. & Bird, A. CpG islands and the regulation of transcription. Genes & Development 25 (10), 1010–1022 (2011). URL https://www.ncbi.nlm.nih.gov/pmc/articles/PMC3093116/. https://doi.org/10.1101/gad.2037511.

[61] Field Guide to Methylation Methods (2016). URL https://www.illumina.com/content/dam/illumina-marketing/documents/products/other/fieldguide_methylation.pdf.

[62] Bird, A. DNA methylation patterns and epigenetic memory. Genes & Development 16 (1), 6–21 (2002). https://doi.org/10.1101/gad.947102.

[63] Heil, B. J. et al. Reproducibility standards for machine learning in the life sciences. Nature Methods 18 (10), 1132–1135 (2021). URL https://www.nature.com/articles/s41592-021-01256-7. https://doi.org/10.1038/s41592-021-01256-7.

[64] Beaulieu-Jones, B. K. & Greene, C. S. Reproducibility of computational workflows is automated using continuous analysis. Nature biotechnology 35 (4), 342–346 (2017). URL https://www.ncbi.nlm.nih.gov/pmc/articles/PMC6103790/. https://doi.org/10.1038/nbt.3780.

[65] Leinonen, R., Sugawara, H., Shumway, M. & International Nucleotide Sequence Database Collaboration. The sequence read archive. Nucleic Acids Research 39 (Database issue) (2011). https://doi.org/10.1093/nar/gkq1019.

[66] Noble, A. J. et al. A validation of Illumina EPIC array system with bisulfite-based amplicon sequencing. PeerJ 9, e10762 (2021). https://doi.org/10.7717/peerj.10762.

[67] Wang, T. et al. A systematic study of normalization methods for Infinium 450K methylation data using whole-genome bisulfite sequencing data. Epigenetics: official journal of the DNA Methylation Society 10 (7), 662–669 (2015). https://doi.org/10.1080/15592294.2015.1057384.

[68] Fischer, B., Smith, M. & Pau, G. rhdf5: R Interface to HDF5 (2020). URL https://github.com/grimbough/rhdf5. R package version 2.35.0.

[69] Pagés, H. HDF5Array: HDF5 backend for DelayedArray objects (2021). URL https://bioconductor.org/packages/HDF5Array. R package version 1.20.0.

[70] Pagés, H., Hickey, P. & Lun, A. DelayedArray: A unified framework for working transparently with on-disk and in-memory array-like datasets (2021). URL https://bioconductor.org/packages/DelayedArray. R package version 0.18.0.

[71] Lowe, R. Marmal-aid - a database for Infinium HumanMethylation450. BMC Bioinformatics 14 (1), 359 (2013). URL https://doi.org/10.1186/1471-2105-14-359. https://doi.org/10.1186/1471-2105-14-359.

[72] Bernstein, M. N., Doan, A., Dewey, C. N. & Wren, J. MetaSRA: normalized human sample-specific metadata for the Sequence Read Archive. Bioinformatics 33 (18), 2914–2923 (2017). URL https://academic.oup.com/bioinformatics/article/33/18/2914/3848915. https://doi.org/10.1093/bioinformatics/btx334.

[73] Fortin, J.-P., Triche, T. J. & Hansen, K. D. Preprocessing, normalization and integration of the illumina humanmethylationepic array with minfi. Bioinformatics 33 (4) (2017). https://doi.org/10.1093/bioinformatics/btw691.

[74] Ritchie, M. E. et al. limma powers differential expression analyses for RNA-sequencing and microarray studies. Nucleic Acids Research 43 (7), e47 (2015). https://doi.org/10.1093/nar/gkv007.

[75] Williams, R. A new algorithm for optimal 2-constraint satisfaction and its implications. Theoretical Computer Science 348 (2), 357–365 (2005). URL https://www.sciencedirect.com/science/article/pii/S0304397505005438. https://doi.org/10.1016/j.tcs.2005.09.023.

[76] Kane, D. M. & Nelson, J. Sparser johnson-lindenstrauss transforms. Journal of the ACM (JACM) 61 (1), 1–23 (2014).

[77] Aumüller, M., Bernhardsson, E. & Faithfull, A. ANN-Benchmarks: A Benchmarking Tool for Approximate Nearest Neighbor Algorithms. 1807.05614 [cs] (2018). URL http://arxiv.org/abs/1807.05614. ArXiv: 1807.05614.

[78] Harris, C. R. et al. Array programming with NumPy. Nature 585 (7825), 357–362 (2020). URL https://doi.org/10.1038/s41586-020-2649-2. https://doi.org/10.1038/s41586-020-2649-2.

[79] Weinberger, K., Dasgupta, A., Attenberg, J., Langford, J. & Smola, A. Feature Hashing for Large Scale Multitask Learning. 0902.2206 [cs] (2010). URL http://arxiv.org/abs/0902.2206. ArXiv: 0902.2206.

[80] Chen, Y.-a. et al. Discovery of cross-reactive probes and polymorphic CpGs in the Illumina Infinium HumanMethylation450 microarray. Epigenetics 8 (2), 203–209 (2013). URL http://www.ncbi.nlm.nih.gov/pmc/articles/PMC3592906/. https://doi.org/10.4161/epi.23470.

[81] Leek, J. T. et al. sva: Surrogate Variable Analysis (2021). R package version 3.40.0.

[82] R Core Team. R: A Language and Environment for Statistical Computing. R Foundation for Statistical Computing, Vienna, Austria (2021). URL https://www.R-project.org/.

[83] Python Core Team. Python: A dynamic, open source programming language. Python Software Foundation (2019). URL https://www.python.org/. Python version 3.7.

[84] Wickham, H. ggplot2: Elegant Graphics for Data Analysis (Springer-Verlag New York, 2016). URL https://ggplot2.tidyverse.org.

[85] Gehlenborg, N. UpSetR: A More Scalable Alternative to Venn and Euler Diagrams for Visualizing Intersecting Sets (2019). URL https://CRAN.R-project.org/package=UpSetR. R package version 1.4.0.

[86] Benjamini, Y. & Hochberg, Y. Controlling the false discovery rate: A practical and powerful approach to multiple testing. JSRRB 57, 289–300 (1995).

